# A guide to performing Polygenic Risk Score analyses

**DOI:** 10.1101/416545

**Authors:** Shing Wan Choi, Timothy Shin Heng Mak, Paul F. O’Reilly

## Abstract

The application of polygenic risk scores (PRS) has become routine across genetic research. Among a range of applications, PRS are exploited to assess shared aetiology between phenotypes, to evaluate the predictive power of genetic data for use in clinical settings, and as part of experimental studies in which, for example, experiments are performed on individuals, or their biological samples (eg. tissues, cells), at the tails of the PRS distribution and contrasted. As GWAS sample sizes increase and PRS become more powerful, they are set to play a key role in personalised medicine. However, despite the growing application and importance of PRS, there are limited guidelines for performing PRS analyses, which can lead to inconsistency between studies and misinterpretation of results. Here we provide detailed guidelines for performing polygenic risk score analyses relevant to different methods for their calculation, outlining standard quality control steps and offering recommendations for best-practice. We also discuss different methods for the calculation of PRS, common misconceptions regarding the interpretation of results and future challenges.

---

Genome-wide association studies (GWAS) have identified a large number of genetic variants, typically single nucleotide polymorphisms (SNP), associated with a wide range of complex traits [1–3]. However, the majority of these variants have a small effect and typically correspond to a small fraction of truly associated variants, meaning that they have limited predictive power [4–6]. Using a linear mixed model in the Genome-wide Complex Trait Analysis software (GCTA) [7], Yang et al (2010) demonstrated that much of the heritability of height can be explained by evaluating the effects of all SNPs simultaneously [6]. Subsequently, statistical techniques such as LD score regression (LDSC) [8,9] and the polygenic risk score (PRS) method [4,10] have also aggregated the effects of variants across the genome to estimate heritability, to infer genetic overlap between traits and to predict phenotypes based on genetic profile or that of other phenotypes [4,5,8–10].

While GCTA, LDSC and PRS can all be exploited to infer heritability and shared aetiology among complex traits, PRS is the only approach that provides an estimate of genetic propensity to a trait at the individual-level. In the standard approach [4,11–13], polygenic risk scores are calculated by computing the sum of risk alleles corresponding to a phenotype of interest in each individual, weighted by the effect size estimate of the most powerful GWAS on the phenotype. Studies have shown that substantially greater predictive power can usually be achieved by using PRS rather than a small number of genome-wide significant SNPs [11,14,15]. As an individual-level genome-wide genetic proxy of a trait, PRS are suitable for a range of applications. For example, as well as identifying shared aetiology among traits, PRS have been used to test for genome-wide G*E and G*G interactions [15,16], to perform Mendelian Randomisation studies to infer causal relationships, and for patient stratification and sub-phenotyping [14,15,17,18]. Thus, while polygenic scores represent individual genetic predictions of phenotypes, prediction is generally not the end objective, rather these predictions are then typically used for interrogating hypotheses via association testing.

Despite the popularity of PRS analyses, there are minimal guidelines [13] regarding how best to perform PRS analyses, and no existing summaries of the differences and options among the main PRS approaches. Here we provide a guide to performing polygenic risk score analysis, outlining the standard quality control steps required, options for PRS calculation and testing, and interpretation of results. We also outline some of the challenges in PRS analyses and highlight common misconceptions in the interpretation of PRS and their results. We will not perform a comparison of the power of different PRS methods nor provide an overview of PRS applications, since these are available elsewhere [13,19], and instead focus this article on the issues relevant to PRS analyses irrespective of method used or application, so that researchers have a starting point and reference guide for performing polygenic score analyses.

## 1. Introduction to Polygenic Risk Scores

We define polygenic risk scores, or polygenic scores, as a single value estimate of an individual’s propensity to a phenotype, calculated as a sum of their genome-wide genotypes weighted by corresponding genotype effect sizes – potentially scaled or shrunk – from summary statistic GWAS data. The use of summary statistic data for the genotype effect size estimates differentiates polygenic scores from phenotypic prediction approaches that exploit individual-level data only, in which genotype effect sizes are typically estimated in joint models of multiple variants and prediction performed simultaneously, such as via best linear unbiased prediction (BLUP) [20,21] and least absolute shrinkage and selection operator (LASSO) [22,23]. While we note that such methods may offer great promise in performing powerful prediction within large individual-level data sets [22], we limit our focus to polygenic scores specifically, which we believe are likely to have enduring application due to (i) the desire to test specific hypotheses on locally collected small-scale data sets, (ii) data sharing restrictions, (iii) heterogeneity across data sets, (iv) large general population data sets, such as the UK Biobank [24], having relatively few individuals with specific diseases compared to dedicated case/control studies.

Therefore, PRS analyses can be characterized by the two input data sets that they require: i) base (GWAS) data: summary statistics (e.g. betas, *P*-values) of genotype-phenotype associations at genetic variants (hereafter SNPs) genome-wide, and ii) target data: genotypes and phenotype(s) in individuals of the target sample. If the population-level effects of the SNPs were estimated from the GWAS without error, then the PRS could predict the phenotype of individuals in the target data with variance explained equal to the “chip-heritability” (h_snp_^2^) of the trait [25]. However, due to error in the effect size estimates and inevitable differences in the base and target samples, the predictive power of PRS are typically substantially lower than h_snp_^2^ (see Figure 4a) but tend towards h_snp_^2^ as GWAS sample sizes increase.

Important challenges in the construction of PRS are the selection of SNPs for inclusion in the score and what, if any, shrinkage to apply to the GWAS effect size estimates (see Section 3.1). If such parameters are already known, then PRS can be computed directly on the target individual(s). However, when parameters for generating an optimal PRS are unknown, then the target sample can be used for model training, allowing optimisation of model parameters. How to perform this parameter optimisation without producing overfit PRS is discussed in Section 4.4. First, we outline recommended quality control (QC) of the base and target data. In Figure 1, a flow chart summarises the fundamental features of a PRS analysis and reflects the structure of this guide.

**Figure 1:**
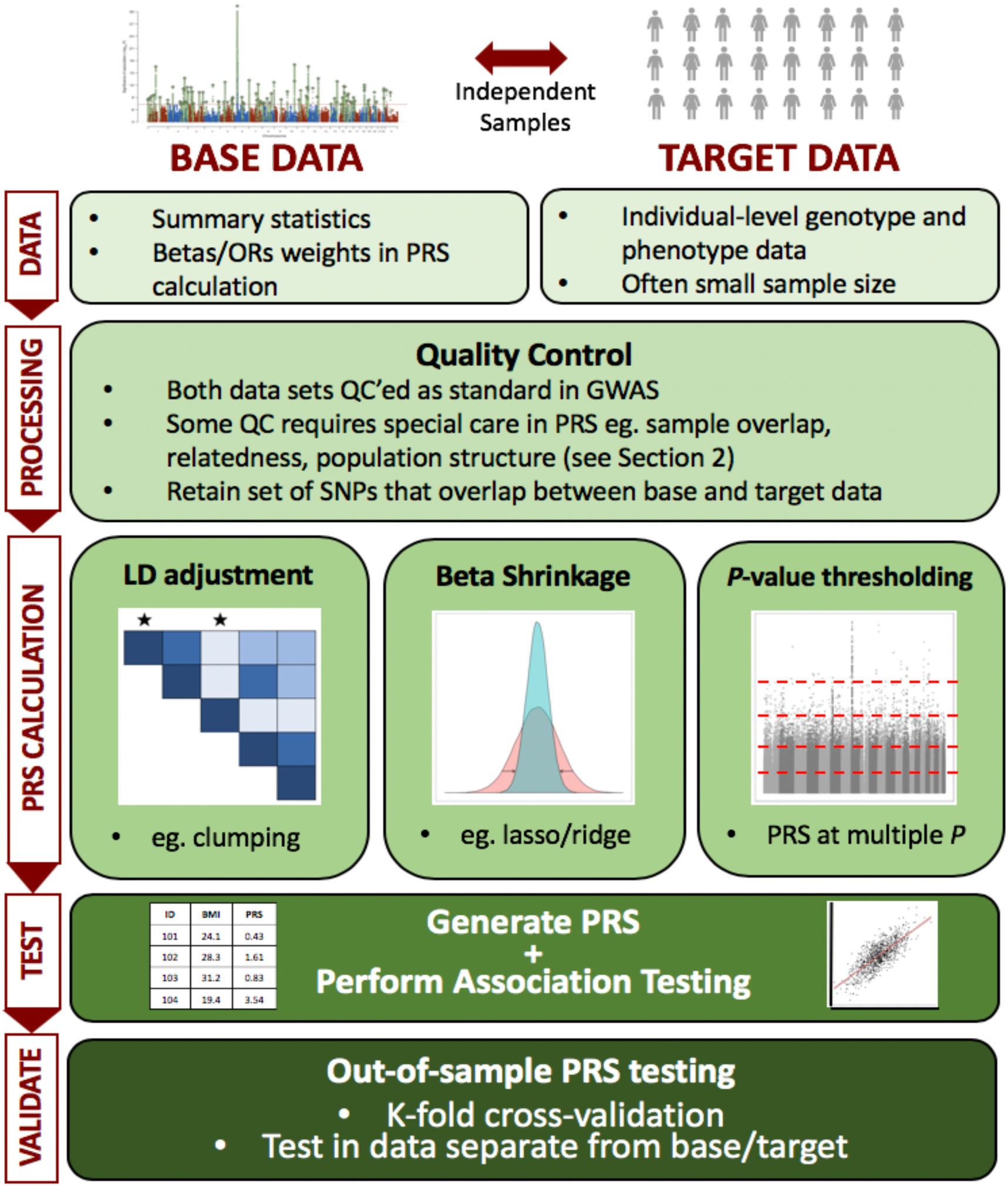
The Polygenic Risk Score (PRS) analysis process. PRS can be defined by their use of base and target data, as in Section 1. Quality control of both data sets is described in Section 2, while the different approaches to calculating PRS – e.g. LD adjustment via clumping, beta shrinkage using lasso regression, *P*-value thresholding – is summarised in Section 3. Issues relating to exploiting PRS for association analyses to test hypotheses, including interpretation of results and avoidance of overfitting to the data, are detailed in Section 4.

## 2. Quality Control of Base and Target data

The power and validity of PRS analyses are dependent on the quality of the base and target data. Therefore, both data sets must be quality controlled to the high standards implemented in GWAS studies, e.g. removing SNPs according to low genotyping rate, minor allele frequency or imputation ‘info score’ and individuals with low genotyping rate (see [26–28]). PLINK is a useful software for performing such quality control (QC) [29,30]. Particular care should be taken over these standard QC procedures since any errors that occur may aggregate across SNPs when PRS are computed. In addition to these standard GWAS QC measures, the following QC issues more specific to PRS analyses need special attention and should act as a checklist for PRS analyses:

### File transfer

Since most base GWAS data are downloaded online, and base/target data transferred internally, one should ensure that files have not been corrupted during transfer, e.g. using md5sum [31]. PRS calculation errors are often due to corrupt files.

### Genome Build

Ensure that the base and target data SNPs have genomic positions assigned on the same genome build [32]. LiftOver [33] is an excellent tool for standardizing genome build across different data sets.

### Effect allele

Some GWAS results files do not make clear which allele is the effect allele and which the non-effect allele. If the incorrect assumption is made in computing the PRS, then the effect of the PRS in the target data will be in the wrong direction, and so to avoid misleading conclusions it is critical that the effect allele from the base (GWAS) data is known.

### Ambiguous SNPs

If the base and target data were generated using different genotyping chips and the chromosome strand (+/-) for either is unknown, then it is not possible to match ambiguous SNPs (i.e. those with complementary alleles, either C/G or A/T) across the data sets, because it will be unknown whether the base and target data are referring to the same allele or not. While allele frequencies can be used to infer which alleles match [34], we recommend removing all ambiguous SNPs since the allele frequencies provided in base GWAS are often those from resources such as the 1000G project, and so aligning alleles according to their frequency could lead to systematic biases in PRS analyses. When there is a non-ambiguous mismatch in allele coding between the data sets, such as A/C in the base and G/T in the target data, then this can be resolved by ‘flipping’ the alleles in the target data to their complementary alleles. Most polygenic score software can perform this flipping automatically.

### Duplicate SNPs

Ensure that there are no duplicated SNPs in either the base or target data since this may cause errors in PRS calculation unless the code/software used specifically checks for duplicated SNPs.

### Sex-check

While sex-check procedures are standard in GWAS QC, they are critical in PRS analyses because errors may generate false-positive associations that are due to sex differences in the target phenotype generated by factors other than autosomal genetics. If the aim is to only model autosomal genetics, then all X and Y chromosome SNPs should be removed from the base and target data to eliminate the possibility of confounding by sex. Proper modelling of the sex chromosomes would improve the predictive power of PRS, but a lack of consensus on how best to analyse the sex chromosomes in GWAS has meant that they have, unfortunately, not generally been considered in PRS studies to date.

### Sample overlap

Sample overlap between the base and target data can result in substantial inflation of the association between the PRS and trait tested in the target data [35] and so must be eliminated either, (1) directly: either removing overlapping samples from the target data, or if this removes most/all target individuals, then in the base data followed by recalculation of the base GWAS, or (2) indirectly: if, and only if, the overlapping samples correspond to the entire target sample, and the GWAS that contributed to the base data is available for use, then the overlap can be eliminated using the analytic solution described in [36]. We expect a correction in more complex scenarios of sample overlap, when these solutions are unavailable, to be an objective of future methods development.

### Relatedness

A high degree of relatedness among individuals between the base and target data can also generate inflation of the association between the PRS and target phenotype. Assuming that the results of the study are intended to reflect those of the general population without close relatedness between the base and target samples, then relatives should be excluded. If genetic data from the relevant base data samples can be accessed, then any closely related individuals (eg. 1^st^/2^nd^ degree relatives) across base and target samples should be removed. If this is not an option, then every effort should be made to select base and target data that are very unlikely to contain highly related individuals.

### Heritability check

A critical factor in the accuracy and predictive power of PRS is the power of the base GWAS data [4], and so to avoid reaching misleading conclusions from the application of PRS we recommend first performing a heritability check of the base GWAS data. We suggest using a software such as LD Score regression [8] or LDAK [37] to estimate chip-heritability from the GWAS summary statistics, and recommend caution in interpretation of PRS analyses that are performed on GWAS with a low chip-heritability estimate (eg. h_snp_^2^ < 0.05).

## 3. The Calculation of Polygenic Risk Scores

Once quality control has been performed on the base and target data, and the data files are formatted appropriately, then the next step is to calculate polygenic risk scores for all individuals in the target sample. There are several options in terms of how PRS are calculated. GWAS are performed on finite samples drawn from particular subsets of the human population, and so the SNP effect size estimates are some combination of true effect and stochastic variation – producing ‘winner’s curse’ among the top-ranking associations – and the estimated effects may not generalise well to different populations (Section 3.4). The aggregation of SNP effects across the genome is also complicated by the correlation among SNPs – ‘Linkage Disequilibrium’ (LD). Thus, key factors in the development of methods for calculating PRS are the potential adjustment of GWAS estimated effect sizes via e.g. shrinkage and incorporation of their uncertainty, (ii) the tailoring of PRS to target populations, and (iii) the task of dealing with LD. We discuss these issues below, and also those relating to the units that PRS values take, the prediction of traits different from the base trait, and multi-trait PRS approaches. Each of these issues should be considered when calculating PRS – though several are automated within specific PRS software – irrespective of application or whether the PRS will be subsequently used for prediction as an end point or for association testing of hypotheses.

### 3.1 Shrinkage of GWAS effect size estimates

Given that SNP effects are estimated with uncertainty and since not all SNPs influence the trait under study, the use of unadjusted effect size estimates of all SNPs could generate poorly estimated PRS with high standard error. To address this, two broad shrinkage strategies have been adopted: i) shrinkage of the effect estimates of all SNPs via standard or tailored statistical techniques, and ii) use of *P*-value selection thresholds as inclusion criteria for SNPs into the score.

(i) PRS methods that perform shrinkage of all SNPs [19,38] generally exploit commonly used statistical shrinkage/regularisation techniques, such as LASSO or ridge regression [19], or Bayesian approaches that perform shrinkage via prior distribution specification [38]. Under different approaches or parameter settings, varying degrees of shrinkage can be achieved: some force most effect estimates to zero or close to zero, some mostly shrink small effects, while others shrink the largest effects most. The most appropriate shrinkage to apply is dependent on the underlying mixture of null and true effect size distributions, which are likely a complex mixture of distributions that vary by trait. Since the optimal shrinkage parameters are unknown *a priori*, PRS prediction is typically optimised across a range of (tuning) parameters (for overfitting issues relating to this, see Section 4.4), which in the case of LDpred, for example, includes a parameter for the fraction of causal variant [38].
(ii) In the *P*-value selection threshold approach, only those SNPs with a GWAS association *P*-value below a certain threshold (eg. *P* < 1×10^−5^) are included in the calculation of the PRS, while all other SNPs are excluded. This approach effectively shrinks all excluded SNPs to an effect size estimate of zero and performs no shrinkage on the effect size estimates of those SNPs included. Since the optimal *P*-value threshold is unknown *a priori*, PRS are calculated over a range of thresholds, association with the target trait tested for each, and the prediction optimised accordingly (see Section 4.4). This process is analogous to tuning parameter optimisation in the formal shrinkage methods. An alternative way to view this approach is as a parsimonious variable selection method, effectively performing forward selection ordered by GWAS *P-*value, involving block-updates of variables (SNPs), with size dependent on the increment between *P*-value thresholds. Thus the ‘optimal threshold’ selected is defined as such only within the context of this forward selection process; a PRS computed from another subset of the SNPs could be more predictive of the target trait, but the number of subsets of SNPs that could be selected is too large to feasibly test given that GWAS are based on millions of SNPs.

### 3.2 Controlling for Linkage Disequilibrium

The association tests in GWAS are typically performed one-SNP-at-a-time, which, combined with the strong correlation structure across the genome, makes identifying the independent genetic effects (or best proxies of these if not genotyped/imputed) extremely challenging. While the power of GWAS can be increased by conditioning on the effects of multiple SNPs simultaneously [39], this requires access to raw data on all samples, so researchers generally need to exploit standard GWAS (one-SNP-at-a-time) summary statistics to compute polygenic scores. There are two main options for approximating the PRS that would have been generated from full conditional GWAS: (i) SNPs are *clumped* so that the retained SNPs are largely independent of each other and thus their effects can be summed, assuming additivity, all SNPs are included and the linkage disequilibrium (LD) between them is accounted for. Usually option (i) is chosen in the ‘standard approach’ to polygenic scoring, involving *P*-value thresholding, while option (ii) is generally favoured in methods that implement traditional shrinkage methods [19,38] (see Table 1). In relation to (i), some researchers, however, prefer to perform the *P*-value thresholding approach without clumping, meaning that the effects of correlated SNPs are summed as though they were independent. While breaking this assumption may lead to minimal losses in some scenarios [19], we recommend performing clumping [13] when non-shrunk effect sizes estimates from GWAS are used because the non-uniform nature of LD across the genome is likely to generate some bias in estimates. The reason why the standard approach, though simple, appears to perform comparably to more sophisticated approaches [19,38] may be due to the clumping process capturing conditionally independent effects well; note that, (i) clumping does not merely thin SNPs by LD at random (like *pruning*) but preferentially selects SNPs most associated with the trait under study, (ii) clumping can retain multiple independent effects in the same genomic region if they exist (it does not simply retain only the most associated SNP in a region). A criticism of clumping, however, is that researchers typically select an arbitrarily chosen correlation threshold [35] for the removal of SNPs in LD, and so while no strategy is without arbitrary features, this may be an area for future development of the classical approach.

**Table 1.** Comparison of different approaches for performing Polygenic Risk Score analyses

| | $P$-value<br>thresholding<br>w/o clumping | *Standard approach:*<br>Clumping +<br>thresholding (C+T) | Penalised<br>Regression | Bayesian<br>Shrinkage |
| --- | --- | --- | --- | --- |
| Shrinkage strategy | $P$-value<br>threshold | $P$-value threshold | LASSO,<br>Elastic Net,<br>penalty<br>parameters | Prior<br>distribution,<br>e.g. fraction of<br>causal SNPs |
| Handling Linkage<br>Disequilibrium | N/A | Clumping | LD matrix is<br>integral to<br>algorithm | Shrink effect<br>sizes with<br>respect to LD |
| Example software | PLINK | PRSice [12] | Lassosum [19] | LDpred [38] |

### 3.3 PRS units

When calculating PRS, the units of the GWAS effect sizes determine the units of the PRS; e.g. if calculating a height PRS using effect sizes from a height GWAS that are reported in centimetres (cm), then the resulting PRS will also be in units of cm. PRS may then be standardised, dividing by the number of SNPs to ensure a similar scale irrespective of number of SNPs included, or standardised to a standard normal distribution. However, the latter discards information that may wish to be retained, since the absolute values of the PRS may be useful in detecting problems with the calculation of the PRS or the sample, identifying outliers, comparing or combining PRS across different samples, or even detecting the effects of natural selection. Negative selection against effect alleles could result in a PRS with a mean negative value due to effect alleles being at lower frequency than non-effect alleles on average, and the opposite for traits under positive selection.

In calculating PRS on a binary (case/control) phenotype, the effect sizes used as weights are typically reported as log Odds Ratios (log(ORs)). Assuming that relative risks on a disease accumulate on a multiplicative rather than additive scale [40], then PRS should be computed as a summation of log(OR)-weighted genotypes. It is important for subsequent interpretation to know which logarithmic scale was used since the PRS will take the same units and will be needed to transform back to an OR scale.

### 3.4 Population structure and global heterogeneity

Population structure is the principal source of confounding in GWAS (post-QC), and thus risk of false-positive findings. Briefly, structure in mating patterns in a population generates structure in genetic variation, correlated most strongly with geographic location, and environmental risk factors can be similarly structured; this creates the potential for associations between many genetic variants and the tested trait that are confounded by e.g. location [41,42]. While this problem is typically addressed in GWAS via adjustment by principal components (PCs) [41] or the use of mixed models [43], population structure poses a potentially greater problem in PRS analyses, because a large number of null variants are typically included in the calculation of PRS and their estimated effects are aggregated. If allele frequencies differ systematically between the base and target data, which may derive from genetic drift or the ascertainment of genotyped variants [44], and if the distributions of environmental risk factors for the trait also differ between the two – both highly likely in most PRS studies – then there is a danger that an association between the PRS and target trait can be generated by differences at null SNPs. Confounding is potentially reintroduced even if the GWAS had controlled for population structure perfectly, because this does not account for correlated differences in allele frequencies and risk factors between the base and target data. When the base and target samples are drawn from the same or genetically similar populations, stringent control for structure in the PRS analysis itself (e.g. including a large number of PCs) should suffice to avoid false-positive findings, but we recommend in general that extreme caution is taken given dramatic differences in PRS distributions observed between populations [44–46]. While these observations do not imply large differences in aetiology across populations – although genuine differences due to variation in the environment, culture and selection pressures are likely to contribute – they do question the reliability of PRS analyses using base and target data from different populations that do not rigorously address the issue of potential confounding from geographic stratification [45]. It is also important to be wary of the fact that highly significant results can be observed due to subtle confounding when exploiting large sample sizes. Note that we use the term ‘population’ here in a statistical sense: problems of population structure are just as relevant within-country given differences in the genetics and environment between individuals in the base and target samples. We expect the issue of the generalisability of PRS across populations to be an active area of methods development in the coming years [46,47].

### 3.5 Predicting Different Traits and exploiting multiple PRS

While PRS are often analysed in scenarios in which the base and target phenotype are the same, many published studies involve a target phenotype different from that on which the PRS is based. These analyses fall into three main categories: (i) optimising target trait prediction using a different but similar (or ‘proxy’) base trait: if there is no large GWAS on the target trait, or it is underpowered compared to a similar trait, then prediction may be improved using a different base trait (e.g. education years to predict cognitive performance [48,49]), (ii) optimising target trait prediction by exploiting multiple PRS based on a range of different traits in a joint model, (iii) testing for evidence of shared aetiology between base and target trait [50]. Applications (i) and (ii) are straightforward in their aetiology-agnostic aim of optimising prediction, achieved by exploiting the fact that a PRS based on one trait is predictive of genetically correlated traits, and that a PRS computed from any base trait is sub-optimal due to the finite size of any GWAS. Application (iii) is inherently more complex because there are different ways of defining and assessing ‘shared aetiology’. Shared aetiology may be due to so-called horizontal pleiotropy (separate direct effects) or vertical pleiotropy (downstream effect) [51] and it is unclear what quantity should be estimated to assess evidence – genetic correlation [9], genetic contribution to phenotypic covariance (co-heritability) [52,53], or a trait-specific measure (eg. where the denominator relates only to the genetic risk of one trait).

While there is active method development in these areas [54–56] at present, the majority of PRS studies use exactly the same approach to PRS analysis whether or not the base and target phenotypes differ [50,56]. However, this is rather unsatisfactory because of the non-uniform genetic sharing between different traits. In PRS analysis, the effect sizes and *P*-values are estimated using the base phenotype, independent of the target phenotype. Thus, a SNP with high effect size and significance in the base GWAS may have no effect on the target phenotype. The standard approach could be adapted so that SNPs are prioritized for inclusion in the PRS according to joint effects on the base and target traits [57], but this has yet to be implemented in any standard software. Other more sophisticated solutions are presently being investigated [55] and other approaches will likely be developed in future, each tailored to specific scientific questions.

## 4. Interpretation and Presentation of Results

If performing individual prediction is the end objective – for example, to make clinical decisions about individual patients – then the most predictive polygenic score method (known at the time) should be applied to the most powerful base sample available on the relevant trait, in order to optimise accuracy of the individual PRS. Little interpretation or presentation of results are required in this setting, and thus Section 4 is devoted to the primary use of PRS in association testing of scientific hypotheses. Once PRS have been calculated, selecting from the options described in Section 3, typically a regression is then performed in the target sample, with the PRS as a predictor of the target phenotype, and covariates included as appropriate.

### 4.1 Association and goodness-of-fit metrics

A typical PRS study involves testing evidence for an association between a PRS and a trait or measuring the extent of the association in the entire or specific strata of the sample. The standard ways of measuring associations in epidemiology, and their related issues, apply here. The association between PRS and outcome can be measured with the standard association or goodness-of-fit metrics, such as the effect size estimate (beta or OR), phenotypic variance explained (R^2^), area under the curve (AUC), and *P*-value corresponding to a null hypothesis of no association. The PRS for many traits are such weak proxies of overall genetic burden (presently) that the phenotypic variance that they explain is often very small (R^2^ < 0.01), although this is not important if the aim is only to establish whether an association exists. However, present evidence on the ubiquity of pleiotropy across the genome [51] indicates that there may be shared aetiology between the vast majority of phenotypes, detectable with sufficient sample size. Thus, establishing the relative extent of associations among a range of traits may be more worthwhile [36,58], and act as a step towards identifying the casual mechanisms underlying these genetic associations [59].

While variance explained (R^2^) is a well-defined concept for continuous trait outcomes, only conceptual proxies of this measure (“pseudo-R^2^”) are available for case/control outcomes. A range of pseudo-R^2^ metrics are used in epidemiology [60,61], with Nagelkerke R^2^ perhaps the most popular. However, Nagelkerke R^2^ suffers from particular bias when the case/control proportion is not reflective of the case population prevalence [60], and so in the context of estimating the genetic contribution to a polygenic disease it may be preferable to estimate the phenotypic variance explained on the liability scale. Intuitively, the R^2^ on the liability scale here estimates the proportion of variance explained by the PRS of a hypothetical normally distributed latent variable that underlies and causes case/control status [60,62]. Heritability is typically estimated on the liability scale for case/control phenotypes [13,60,62]. Lee et al [60] developed a pseudo-R^2^ metric that accounts for an ascertained case/control ratio and is measured on the liability scale. We show that, under simulation, this metric indeed controls for case/control ratios that do not reflect disease prevalence, while Nagelkerke R^2^ does not (Figure 2).

**Figure 2.**
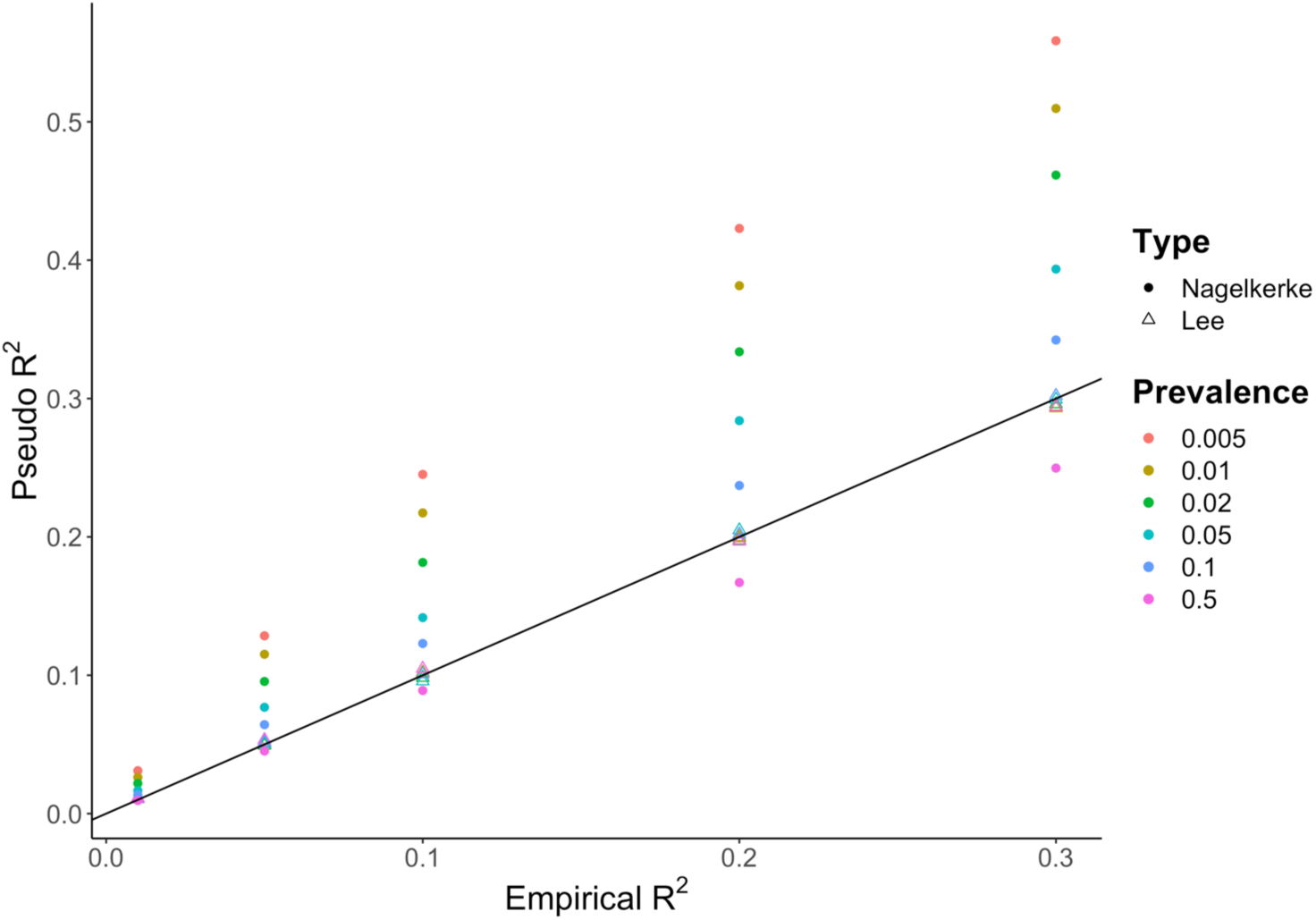
Results from a simulation study comparing Nagelkerke pseudo-R^2^ with the pseudo-R^2^ proposed by Lee et al [59] that incorporates adjustment for the sample case:control ratio. In the simulation, 2,000,000 samples were simulated to have a normally distributed phenotype, generated by a normally distributed predictor (eg. a PRS) explaining a varying fraction of phenotypic variance and a residual error term to model all other effects. Case/control status was then simulated under the liability threshold model according to a specified prevalence. 5,000 cases and 5,000 controls were then randomly selected from the population, and the R^2^ of the original continuous data, estimated by linear regression (Empirical R^2^), was compared to both the Nagelkerke R^2^ (discs) and the Lee R^2^ (triangles) based on the equivalent case/control data by logistic regression.

### 4.2 Graphical representations of results: bar and quantile plots

When the standard approach (C+T) is used, the results of the PRS association testing are often displayed as a bar plot, where each bar corresponds to the result from testing a PRS comprising SNPs with GWAS *P*-value exceeding a given threshold. Typically, a small number of bars are shown, reflecting results at round-figure *P*-value thresholds (5e-8, 1e-5, 1e-3, 0.01, 0.05, 0.1, 0.2, 0.3 etc), and if ‘high-resolution’ scoring [12] is performed then a bar representing the most predictive PRS is included. Usually the Y-axis corresponds to the phenotypic variance explained by the PRS (R^2^ or pseudo-R^2^) and the value over each bar (or its colour) provides the *P*-value of the association between the PRS and target trait. It is important to note that the *P*-value threshold of the most predictive PRS is a function of the effect size distribution, the power of the base (GWAS) and target data, the genetic architecture of the trait, and the fraction of causal variants, and so should not be merely interpreted as reflecting the fraction of causal variants. For example, if the GWAS data are relatively underpowered then the optimal threshold is more likely to be *P* = 1 (all SNPs) even if a small fraction of SNPs is causal (see [4] for details).

While goodness-of-fit measures, such as R^2^, provide a sample-wide summary of the predictive power of a PRS, it can be useful to inspect how trait values vary with increasing PRS or to gauge the elevated disease risk that specific strata of the population may be at according to their PRS. This can be easily visualized using a quantile plot (Figure 3a). Quantile plots in PRS studies are usually constructed as follows [2,63]. The target sample is first separated into strata of increasing PRS. For instance, 20 equally sized quantiles, each comprising 5% of the PRS sample distribution (Figure 3a). The phenotype values of each quantile are then compared to those of the reference quantile (usually the median quantile) one-by-one, with quantile status as predictor of target phenotype (reference quantile coded 0, test quantile coded 1) in a regression. Points on the plot depict the beta or OR (Y-axis), along with bars for their standard errors, corresponding to the regression coefficients of these quantile-status predictors. If covariates are controlled for in the main analysis, then a regression can be performed with target trait as outcome and the covariates as predictors, and the residual trait on the Y-axis instead. Stratification may be performed on unequal strata of PRS (eg. Fig. 3b), in which case these are strata rather than quantile plots. Individuals with high PRS may have low trait values or vice versa, particularly if PRS explain minimal phenotypic variance, and thus the quantiles/strata are not necessarily monotonically increasing.

**Figure 3.**
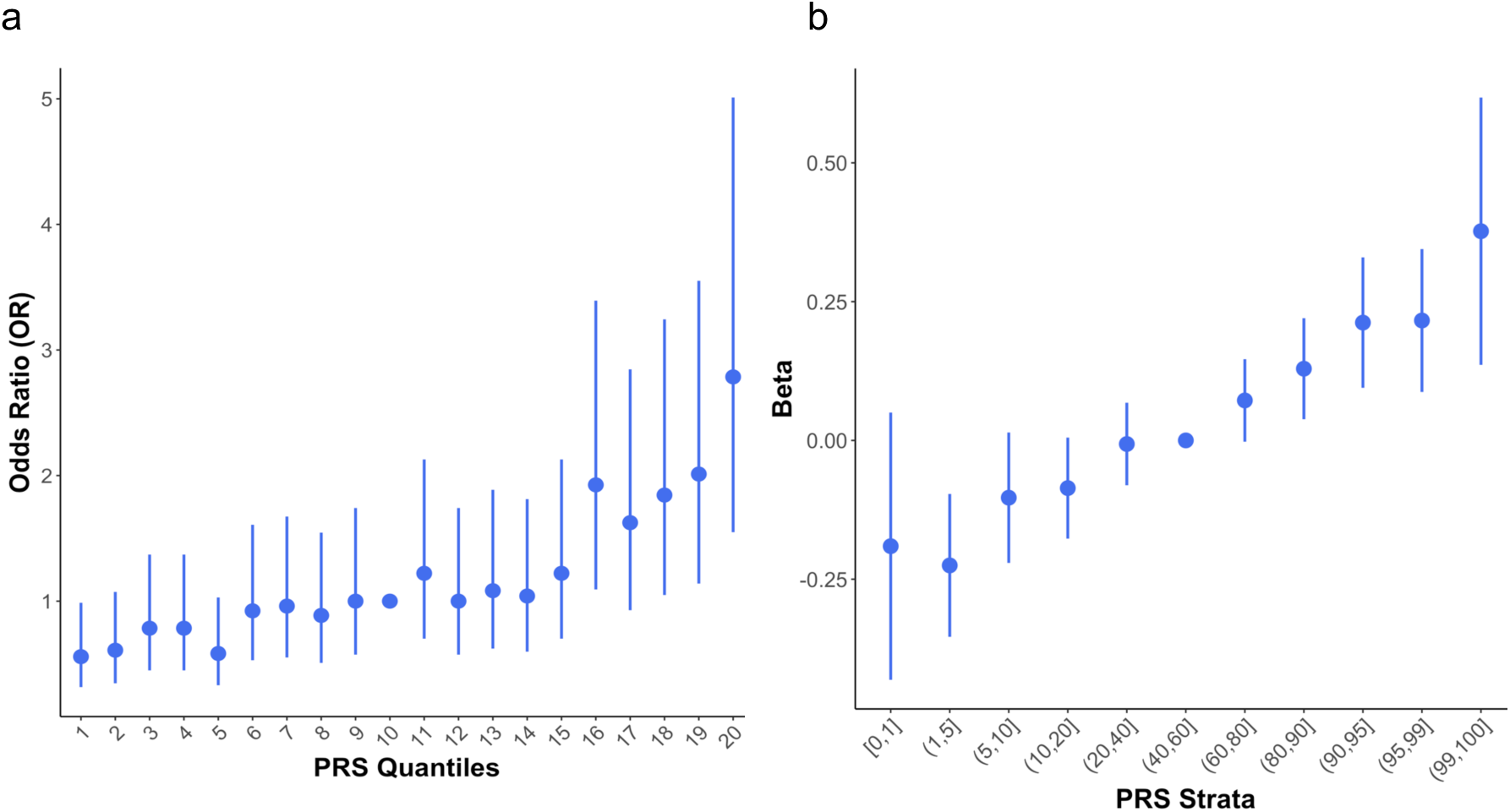
Examples of quantile/strata plots. (a) shows the odds ratios (Y-axis) of a target trait across twenty equal-sized strata of increasing PRS (X-axis) in relation to the 10^th^ strata, while (b) shows a strata plot with eleven unequal strata that highlight the increased or decreased risk among individuals in the top and bottom percentiles of PRS, relative to individuals with PRS in the middle of the distribution (here from 40%-60%).

### 4.3 PRS distribution

Quantile plots corresponding to the same normally distributed phenotype in base and target, should reflect the S-shape of the probit function, and likewise for a binary trait underlain by a normally distributed liability, characterised by the liability threshold model [64]. Thus, inflections of risk at the tails of the PRS distribution [65], or at the top/bottom quantiles, should be interpreted according to this expectation. As for the PRS distribution itself, without respect to association with a target trait, the central limit theorem dictates that if the PRS is based on a sum of independent variables (here SNPs) with identical distributions, then the PRS of a sample should approximate the normal (Gaussian) distribution. Strong violations of these assumptions, such as the use of many correlated SNPs or a sample of heterogenous ancestry (thus SNPs with non-identical genotype distributions), can lead to non-normal PRS distributions. Samples of individuals from disparate worldwide populations may lead to highly non-normal PRS distributions (see Section 3.4), thus inspection of PRS distributions may be informative for problems of population stratification in the target sample not adequately controlled for.

### 4.4 Overfitting in PRS association testing

A common concern in PRS studies that adopt the standard (C+T) approach is whether the use of the most predictive PRS – based on testing at many *P*-value thresholds – overfits to the target data and thus produces inflated results and false conclusions. While such caution is to be encouraged in general, potential overfitting is a normal part of prediction modelling, relevant to the other PRS approaches (Table 1), and there are well-established strategies for optimising power while avoiding overfitting. One strategy that we do not recommend is to perform no optimisation of parameters – e.g. selecting a single arbitrary *P*-value threshold (such as *P <* x10^−8^ or *P* = 1) – because this may lead to serious *underfitting*, which itself can lead to false conclusions.

The gold-standard strategy for guarding against generating overfit prediction models and results is to perform out-of-sample prediction. First, parameters are optimised using a training sample and then the optimised model is tested in a test or validation data set to assess performance. In the PRS setting involving a base and target data, it would be a misconception to believe that out-of-sample prediction has already been performed because polygenic scoring involves two different data sets, when in fact the training is performed on the target data set, meaning that a third data set is required for out-of-sample prediction. In the absence of an independent data set, the target sample can be subdivided into training and validation data sets, and this process can be repeated with different partitions of the sample, e.g. performing 10-fold cross-validation [56,66,67], to obtain more robust model estimates. However, a true out-of-sample, and thus not overfit, assessment of performance can only be achieved via final testing on a sample entirely separate from data used in training.

Without validation data or when the size of the target data makes cross-validation underpowered, an alternative is to generate empirical *P*-values corresponding to the optimised PRS prediction of the target trait, via permutation [12]. While the PRS itself may be overfit, if the objective of the PRS study is association testing of a hypothesis – e.g. H_0_: schizophrenia and rheumatoid arthritis have shared genetic aetiology – rather than for prediction *per se*, then generating empirical *P*-values offers a powerful way to achieve this while maintaining appropriate type 1 error [12]. It is also even possible to generate optimised parameters for PRS when no target data are available [19].

### 4.5 Power and accuracy of PRS: target sample sizes required

In one of the key PRS papers published to date, Dudbridge 2013 [4] provided estimates of the power and predictive accuracy of PRS in different scenarios of data availability and phenotypes. To complement this work, we performed PRS analyses across three traits in the UK Biobank with high (height), medium (Forced Volume Capacity; FVC) and low (hand grip strength) heritability to provide a guide to the approximate performance of PRS association testing on real data with different heritability and different validation sample sizes, when exploiting a large (100k) base GWAS (Figure 4). While this provides only a very limited indication of the performance of PRS analyses, in our experience, researchers in the field often wish to obtain some idea of whether their own (target/validation) data are likely to be sufficiently powered for future analyses or if they need to acquire more data.

**Figure 4.**
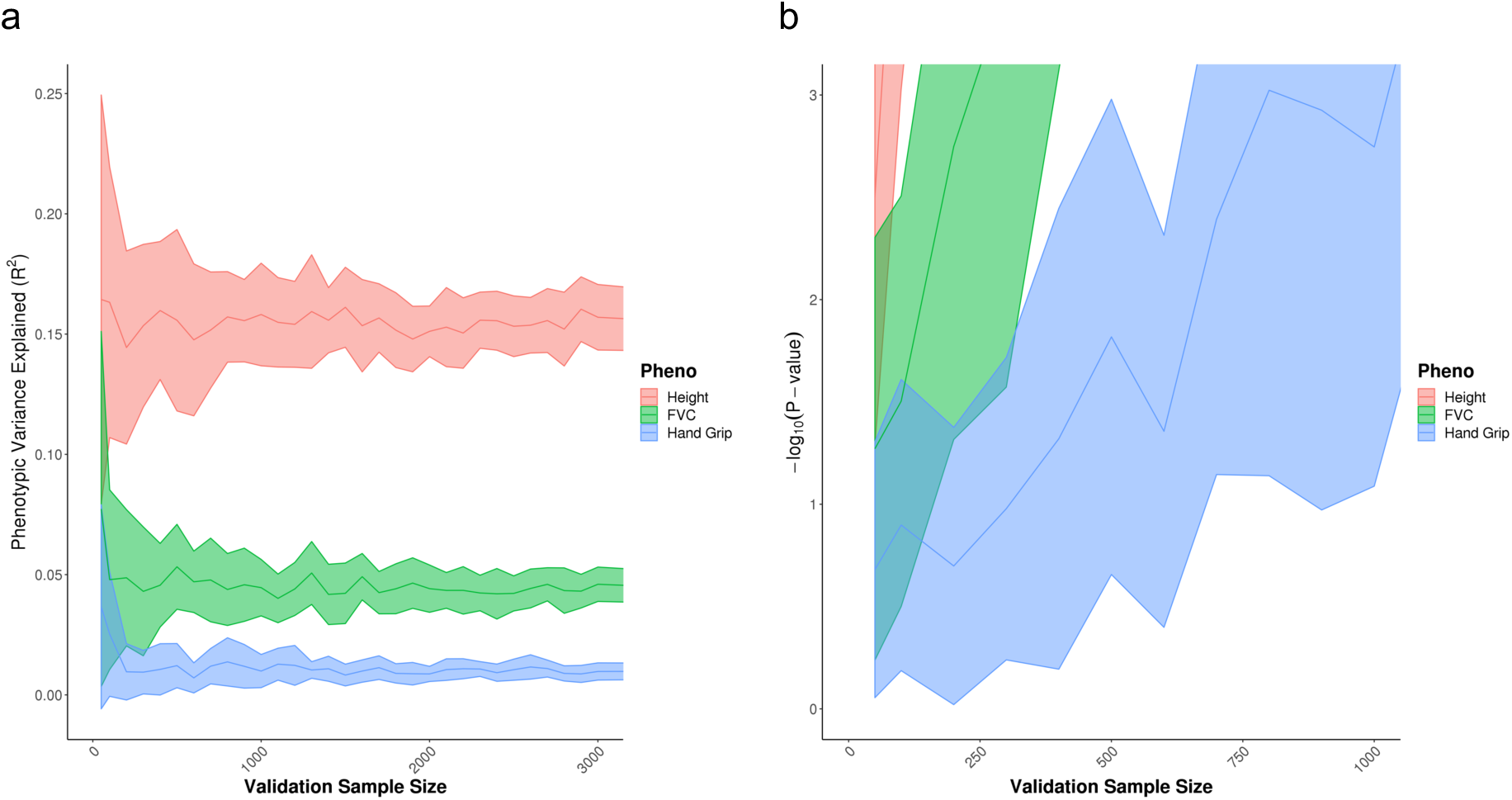
Examples of performance of PRS analyses on real data by validation sample size, according to (a) phenotypic variance explained (R^2^), (b) association *P*-value. UK Biobank data on height (estimated heritability *h*^*2*^ = 0.49 [8]), Forced Volume Capacity (FVC) (estimated heritability *h*^2^ = 0.23 [8]), Hand Grip (estimated heritability *h*^*2*^ = 0.11 [8]), were randomly split into two sets of 100,000 individuals and used as base and target data, while the remaining sample was used as validation data of varying sample sizes, from 50 individuals to 3000 individuals. Each analysis was repeated 5 times with independently selected validation samples. While these results correspond to performance in validation data, the association *P-*values should reflect empirical *P*-values estimated from target data (as described in Section 4.4).

## Conclusions

As GWAS sample sizes increase, the application of Polygenic Risk Scores is likely to play a central role in the future of biomedical studies and personalised medicine. However, the efficacy of their use will depend on the continued development of methods that exploit them, their proper analysis and appropriate interpretation, and an understanding of their strengths and limitations.

## Acknowledgements

We thank the participants in the UK Biobank and the scientists involved in the construction of this resource. This research has been conducted using the UK Biobank Resource under application 18177 (Dr O’Reilly). We thank Jonathan Coleman and Kylie Glanville for help in management of the UK Biobank resource at King’s College London, and we thank Jack Euesden, Tom Bond, Gerome Breen, Cathryn Lewis and Pak Sham for helpful discussions. PFO receives funding from the UK Medical Research Council (MR/N015746/1). SWC is funded from the UK Medical Research Council (MR/N015746/1). This report represents independent research (part)-funded by the National Institute for Health Research (NIHR) Biomedical Research Centre at South London and Maudsley NHS Foundation Trust and King’s College London. The views expressed are those of the authors and not necessarily those of the NHS, the NIHR, or the Department of Health.

## Author contributions

SWC and PFO conceived and prepared the manuscript. All authors discussed its contents. SWC performed all statistical analyses. PFO drafted the manuscript, with critical feedback from SWC and TM.

